# Chemotaxis microsimulation: On the gain in nutrient uptake and bacterial cell division with chemotaxis mechanism

**DOI:** 10.1101/102400

**Authors:** Soutick Saha, C. K. Sruthi, Meher K. Prakash

## Abstract

Bacterial swimming alternates between straight runs for several seconds and tumbles into random directions. Chemotactic bacteria remember nutrient sensing history, change tumble frequency to move toward nutrients. A question that has not been addressed is the significance of the nutrition gain and multiplication of bacterial population with chemotaxis mechanism. To quantify these effects, we introduce a microsimulation model, which seamlessly integrates detailed observations and assumptions about single bacterial tumbles, noisy sensing and nutrient uptake for studying up to a few millions of them in a population. We use the model to simulate absorption of nutrients from lysis and agar plates. Contrary to an intuitive feeling that chemotaxis could be useful under nutrient starvation, we see a significant effect only under nutrient rich conditions where bacteria with chemotaxis outgrow their non-chemotactic counterparts by hundreds of times. The model offers the flexibility to study the consequences of newer assumptions, and experimental conditions.

**Author Summary:** Chemotaxis is a mechanism that helps bacteria navigate towards nutrients. Several aspects of the mechanism have been well studied over the past 50 years. As most bacterial mechanisms are helpful evolutionarily to survive and to multiply, it would be a natural question to ask how much this swim helps bacteria to gain nutrition and consequently to multiply. However to our knowledge this question has not been asked. We develop a model that integrates bacterial motion, with sensing and nutrient uptake and show that only under nutrient rich conditions this mechanism helps.

## Introduction

Bacteria swim toward nutrient source. This movement, called chemotaxis, is guided by nutrient concentration gradients.^1^ In the 1960’s, ring-like patterns formed by *E. coli* in petri-dishes were interpreted as due to a combination of two effects-depletion of the nutrients by bacterial aggregates at the center and a drift towards higher nutrient concentrations.^2,3^ These studies were followed by other population level experiments in semi-solid media which showed the formation of different growth patterns.^4,5^ Pioneering microscopic studies by Berg and others on single bacterium revealed several interesting aspects of the random walk: runs in straight lines followed by sudden tumbles that randomly change the direction,^6^ nutrient memory that *E. coli* holds to bias its walk,^7^ and counter-clockwise and clockwise rotation of flagellar motors that propel the runs and tumbles respectively.^1,8–10^ Concentration-jump experiments on *Salmonella* ^11^ and nutrient release near tethered *E. coli* ^7^ established the bias in run-time lengths with nutrient gradients. The latter study even extracted a form of the memory kernel which describes how the temporal memory of the nutrients encountered by the bacterium biases its run-time lengths.^7^ Further studies have shed light on the molecular details of flagellar motors that propel bacterial chemotaxis,^12–15^ receptor occupancy responsible for the nutrient memory,^16^ signal transduction,^17,18^ and nutrient uptake pathways^19–21^ in chemotaxis.

Mathematical models have been used at different levels of coarse-graining to address the different experimentally relevant questions: population level modeling of bacterial pattern formation in nutrient rich agar gels,^22–24^ biased random-walk and distributions of run, tumble times of individual bacterium,^25^ and robustness of signal transduction pathways,^16,26,27^ as well as strategies for nutrient acquisition, nutrient sensing limits, and consequences for the chemotactic response.^10,28,29^ Further studies have focused on game theoretical strategies^30^ for nutrient acquisition, as well as redistribution of dissolved organic matter in the oceanic environment.^31–34^ Most of these models catered either to the individual bacterium or to the population. The compromise between details and size of the population left gaps in between. Active walker models,^35^ partly tried to bridge this gap. In these, the walkers actively perturb the landscape which defines their movement (in this case bacteria consume nutrients, which eventually affects their trajectory) tracked individual clusters of about 10^3^ bacteria. The focus however was restricted to coarse grained active walkers and was used to study on bacterial pattern formation in semi-solid media, and not specifically on the details of chemotactic movement. Other models studied pattern formation in a fixed population (10^3^) of quorum sensing bacteria under confinement. While the model considered the details of bacterial runs and tumbles as well as the diffusion of quorum sensing molecules, nutrient absorption and cell division were out of the scope of that work.^36^

Fundamental to chemotaxis is a purpose to gain nutrition which possibly helps bacterial growth,^37^ however the relation between the chemotactic movement strategy and cell-division has not been modeled. Bacterial cell division depends neither on the local concentration of the nutrient or an average absorption in the population. Instead each bacterial cell divides using the nutrient it uptakes. Most population level models did not explicitly account for the functional form of the memory that was carefully derived from experiments^7^ with an exception of a few attempts to relate the nutrient memory kernel to drift in the diffusive equation.^38^ Different strategies for biasing the random-walk were proposed by Koshland^11^ and Berg^7^ for *Salmonella* and *E. Coli* respectively. The effectiveness of these different biasing strategies, which we call *minmax* and *min* tumble rate strategies (described in Methods section), were not compared in any model. Although nutrient absorption by bacteria has been described using Michaelis-Menten kinetics, studies that focused on noisy sensing and nutrient acquisition strategies^30^ assumed a perfect collision-absorption condition for the nutrients and over estimated the absorption. Though the theoretical limits of noisy nutrient sensing under low linear concentration gradient has been studied,^28,29^ it has not been integrated into simulations where nutrient diffusion is studied to model experimental conditions.

Addressing these questions requires incorporation of as many details as possible into the model, which may be compromised if one begins with the population level equations. In order to address these gaps which arise between the microscopic level assumptions about the bacteria and the population level observations, we introduce a microsimulation model. Microsimulation or agent-based models^39^ use a set of rules for how each agent responds to environment or other agents. By simulating several interacting agents, the model aims to gain insights into the emergent behavior at the population level. The models are commonly used in health, finance and traffic modeling. The formalism allows an easy implementation of the detailed agent-level assumptions and population level observations. We illustrate the method by simulating in a very detailed way a single bacterium as well as a colony of 10^5^ *E. Coli* growing in a glucose rich medium.

## Methods

**Memory kernel**: In our model, each bacterium performs random walk with runs at an average speed of *v*_0_ = 30 *µm/s*,^8^ and tumbles with a rate,^10,30^

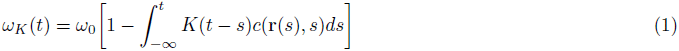

where *c*(**r**(*s*), *s*) is the chemical concentration sensed at a past time *s*,

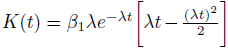

is the memory kernel^7^, and 
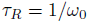
 average run time in the absence of a concentration gradient. Although *β*_1_ = 0.67, *λ*= 2.08*s* ^−1^,^30^ have been mentioned up to a scaling factor, to the best of our knowledge the scaling factor has not been mentioned. We estimate for the scaling factor using the information that the chemotactic response is seen above micromolar concentrations^10^ as such the integral in equation 1 becomes 
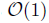
 when 
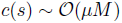
 concentration, where 
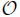
 is used to signify of the order of.

**Uptake**: *E. coli* uptakes using passive and active transport mechanisms.^40^ Glucose is primarily transported in *E. coli* using phosphotransferase system (PTS). Mannose permease is also used to transport glucose with low affinity (at high concentration).^41,42^ Glucose uptake by E. coli follows Michaelis-Menten kinetics. There are three pathways in *E. coli* meant for uptaking glucose in low (< 0.1 *μM*), intermediate (0.1*μM* - 300 *μM*) and high chemical concentrations (> 0.3*mM*) with 0.2 *μM*(*K*_*M*1_), 4 *μM*(*K*_*M*2_) and 1*mM(K_M3_*) as respective half saturation constants^41^ and maximum uptake rate (*V_max_*) of glucose 0.19*fg/cell/s* (Supporting Information).^43^ We assume that in *E. coli*, all three uptake pathways are simultaneously active. The instantaneous uptake rate for the *i^th^* bacterium is:

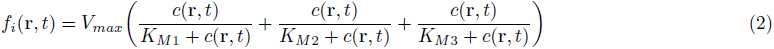

We make a comparison with the perfectly absorbing collision model, assuming the bacterium to be a sphere of radius *a(μm)* and that it perfectly uptakes all nutrient molecules which collide with it, as is assumed in the literature. Nutrients at an average chemical concentration 
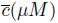
 with diffusivity *D*(*μm*^2^*/s*), have 
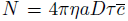
 encounters with the bacterium in time *τ*(*s*), where *η* = 602.3*μm*^-3^*μM*^-1^ is the numerical conversion factor ^28^.

**Mass doubling and cell division**: A single *E. coli* needs *m_D_*= 3 ×10^9^ molecules or 900 fg of glucose in its lifetime, two-thirds for biomass and a third for ATP production^44^. In our model we assume that an *E. coli* divides into two equal halves once it acquires 900 fg of glucose. Until the next tumble event the two daughters have the same position and velocity as their parent at the time of division. If *f_i_*(**r**(*t*), *t*) is the instantaneous total glucose absorption by the *i^th^* bacteria and 
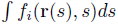
 indicates the cumulative absorption since its birth along the trajectory. Then at time time *t* + *dt*, the total number of bacteria is given by

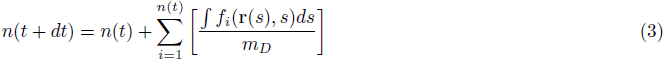

where *dt* is the length of a time step in our simulations, [*q*] is the box function which gives the integral portion of *q* and *n*(0) = *n*_0_ is the size of the initial bacterial population.

**Biasing strategies considered**: Four different memory-dependent biasing strategies were considered in our analysis. The tumble rates *ω* are chosen selectively toward or away from the nutrient after making a comparison between *ω*_0_ and *ω_K_*. The strategies are dubbed as *minmax* tumble rate (*ω* = *ω*_k_) *fixed (ω* = *ω*_0_)^45^ *min* (*ω* = *min*(*ω*_K_, *ω*_0_)) and *max* (*ω* = *max*(*ω*_K_, *ω*_0_)) tumble rate strategies.

**Nutrient diffusion**: The chemical concentration *c*(**r**, *t*) is obtained by combining the diffusion of nutrient with a constant *D* (600 *μm^2^/s* for glucose) and nutrient depletion due to uptake using:

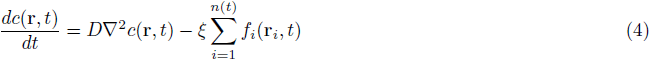

*f_i_*(**r**, *t*) is the nutrient absorbed by *i^th^* bacterium at position **r** and time *t*. **r**_*i*_ is the position of the *i^th^* bacteria which evolves as **r**_*i*_(*t* + *dt*) = **r**_*i*_(*t*) + v_*i*_(*t*)*dt* and *n(t)* be the total bacteria in the population at time *t* and ξ = 1/*dV* * *M_glucose_* where *dV* is the volume of the discretized cells in the numerical solution and *M_glucose_* is the molecular weight of glucose.

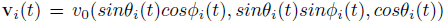

is the three dimensional velocity vector at time *t* defined by the angles *θ_i_(t), ø_i_(t)*. If *ω (t)* is the rate of tumbling according to any of the strategies discussed above, then 
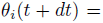
 
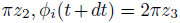
 for *z*_1_ > *w*(*t*)*dt* and

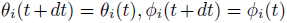
 for *z*_1_ > *ω*(*t*)*dt* where *z*_1_, *z*_2_ and *z*_3_ are three uniform random numbers between 0 and 1. The details of numerical implementation are discussed in Supplementary information.

**Noise and Sensing:** *E. coli* senses the nutrients in its vicinity using receptors of various kinds which are clustered as trimers at the cell poles.^46^ Major receptors include TaR and TsR for sensing aspartate and serine respectively and is abundant in an *E. coli*. Minor receptors like Tap and Trg for sensing dipeptides, ribose and galactose are also present in smaller amounts.^47^ We also assume that the *E. coli* of radius *a(μm)* senses average chemical concentration 
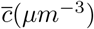
 of molecules with diffusivity *D(μm^2^/s)* in time *τ(s)* with minimum relative error 
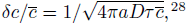

^28^ where *δc* is the absolute error in sensing the chemical concentration. To account for the fluctuations in concentration sensing, the actual chemical concentration sensed by an individual *E. coli* is obtained from a gaussian probability distribution with mean 
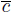

and standard deviation *δc*. The concentration sensed is considered to be zero if the concentration obtained from gaussian distribution is negative, which mostly happens for average concentration less than 1 nM (Figure 4A).

## Results and discussions

### Microsimulation model for chemotactic behavior

There has been a gap in the chemotaxis literature with several interesting microscopic details which are available but not included in the modeling of chemotactic behavior. For example, despite continued research on chemotaxis driven by favorable nutrient gradient, to our knowledge, the nutrient uptake by considering the appropriate absorption kinetics has not been quantitatively assessed. The goal of our chemotaxis microsimulation model (Figure 1) is to be precise with the assumptions at a microscopic level and to make predictions at the population level. Individual bacterial position and nutrient absorption history are completely tracked in our model. Nutrient sensing history, is used along with the experimentally observed memory kernel to bias the walk. The nutrient absorption history modeled using Michaelis-Menten kinetics is used to follow the cell growth and division. The noise in sensing is always explicitly included, although it matters at only extremely low nutrient concentrations (below 1 nM). The microsimulation model is simple and flexible enough that modifications or addition of new details about the physics of the bacterium, such as quorum sensing can be made easily. The largest starting size of the bacterial colony used in our microsimulation is 10^5^ bacterial cells, which was simulated 8 hours of lab experiment to reach a maximum population size of 2.5 × 10^6^ bacteria. To our knowledge this is the largest simulation of a bacterial colony where every individual bacterium is tracked. The computational resources required for this simulation were about 3 GB RAM and 48 hours on a single processor. Simulating a system ten times larger requires ten times larger memory and time and we did not pursue it. Trivial parallelization of the bacteria on multiple processors is not possible since they are connected by the nutrient distribution. Despite this limitation, the microsimulation was useful to address several questions discussed below.

**Figure 1:**
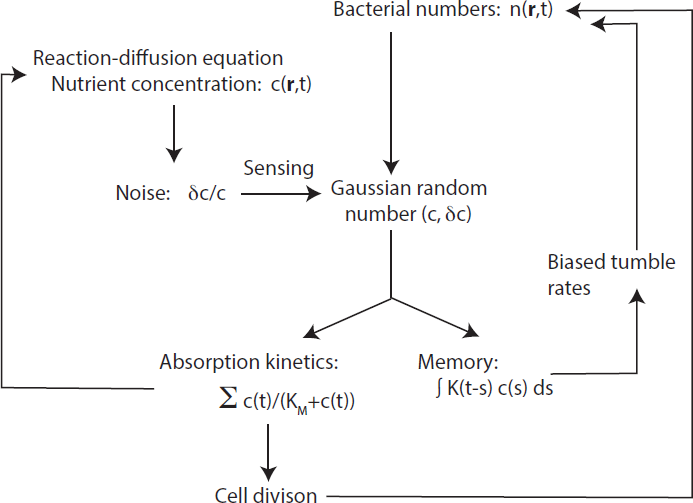
Schematic diagram of microsimulation methodology

### Single bacterium near lysis

#### Uptake from dissolved nutrients

We first analysed the nutrient gain of a single bacterium in a non-equilibrium situation where the nutrients are suddenly released away from the bacterium. One such occurrence in oceanic conditions could be the release of nutrients from phytoplankton lysis. We modeled this event with the burst of a 100 *µ*m radius sphere with 1000 *µ*M glucose concentration. A typical trajectory traced by a chemotactic (*minmax* tumble rate) bacterium is illustrated by considering a bacterium starting at 750 *µm* from nutrient release (Figure 2). We compare the two cases where the background nutrient concentrations are *C_B_* = 0*µ M* and *C_B_* = 40*µM*^48^. The overall nutrient gained for two conditions of the bacterium moving randomly or biased by nutrient gradients using the *fixed* or *minmax* tumble rate strategies are shown in Figure3. Surprisingly a single chemotactic bacterium does not gain much from nutrient uptake under lysis conditions compared to the non-chemotactic bacteria (*fixed* tumble rate) (Figure 3). Infact, under the conditions studied, bacterium gains its nutrient requirement of 900 fg from the dissolved nutrients (*C_B_* = 40*μM*). No significant uptake gain was noticed for lysis events for different distances of the single bacterium from the food source (data not shown). Similarly, no nutrient uptake gain was also noticed for simulating a colony of bacteria near the lysis event (Figure S13) (Supplementary information).

**Figure 2:**
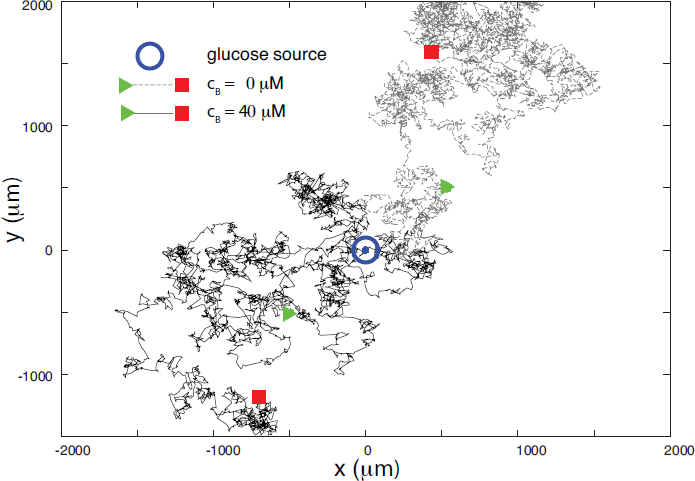
X-Y projection of trajectory of a single *E. coli* near a lysis event in media with a background nutrient concentration of 0 *μ*M (gray lines) and 40 *μ*M (black lines). Lysis event is assumed to be from the burst of a cell with 1000 *μ*M concentration and 100 *μ*m radius and bacteria move with *minmax* tumble rate. Considering the nutrient concentrations involved and the noise in sensing, bacterium ends up drifting away from the source of the lysis burst.

**Figure 3:**
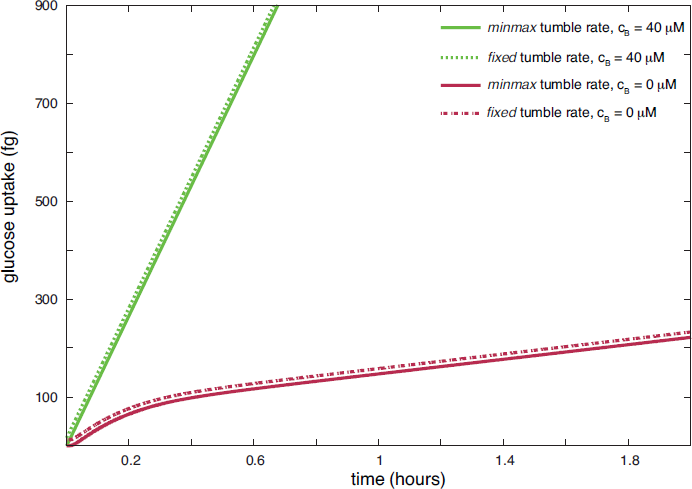
A comparison of nutrient gained by bacteria when swimming with *minmax* tumble rates (solid lines) and a random walk scenario (dotted lines, no concentration-gradient dependent bias) near the lysis event discussed above in Figure 2. Bacteria do not gain enough nutrition from lysis burst, and it comes significantly by absorbing from the uniform background. The dotted lines were shifted up uniformly by 10 fg for clarity.

#### Role of noisy sensing

The role of noise and sensitivity limits have been extensively discussed in the literature.^28, 29^ The minimum relative noise sensed by an *E. coli* of radius *a(µm)* in time *τ(s)* and swimming in an average concentration 
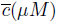
 and the nurients of the medium having diffusivity *D(µm^2^/s)* and conversion factor *η= 602.3 µm ^-3^ µM ^-1^* is given by 
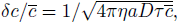

^28^ which is shown in Figure 4A for different concentrations. The deviations of the number of encounters relative to its average thus drops significantly above a concentration of 10 nM. In our microsimulation, we always include the effect of noise, whether or not it is relevant. In every instance of simulation, we generate a gaussian random number using a mean c and standard deviation *δc* and use it for estimating the sensitivity of detection. Also shown in Figure 4A are the instantaneous rate (*f_i_*(**r**, *t*)) of absorption at a given concentration, and the time required for 900 fg absorption assuming this instantaneous rate. The nutrient experienced above background by a single bacterium along the trajectory (Figure2) is shown in Figure 4B. While the noise limits in sensing are interesting, in a background concentration of 40 *μ*M, this noise limit becomes irrelevant. As illustrated by combining all the three graphs in Figure 4A. Interestingly the limits where noise becomes significant are where the nutrient concentration is too low to have a reasonable uptake in several hours.

**Figure 4:**
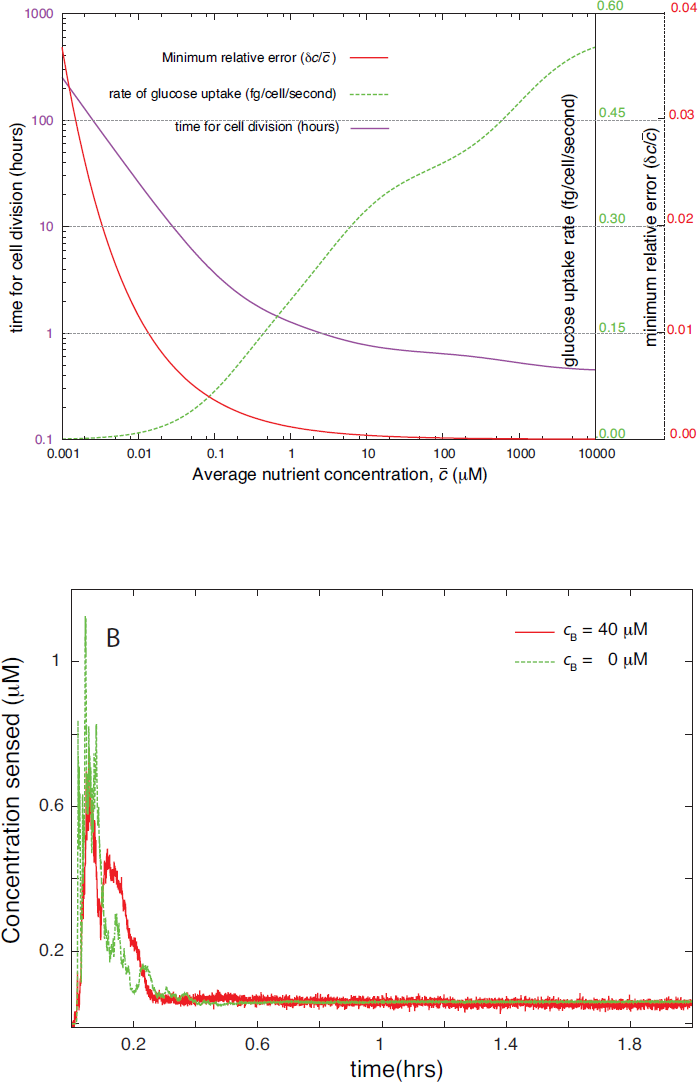
(A) Minimum relative error in nutrient sensing at a given concentration of the nutrients(red line), rate of glucose uptake by a single cell (green line), and time for cell division after absorption of 900 fg of glucose (purple line). (B) The concentration experienced by the bacterium near the lysis event above background in Figure 2.

### *E. Coli* in culture conditions

#### Effect of memory on swarm ring formation

A microsimulation was performed to mimic the growth of a 10^5^ bacterial colony placed at the center of a 40*mm* × 40*mm* × 0.125*mm* petridish with a 100 *μM* uniform glucose concentration^3^. The evolution of the bacterial colony after 6 hours is shown in Figure 5. The simulation with *minmax* tumble strategy replicates the experimentally observed ring formation in Figure 5A because of the nutrient depletion at the center, migration of the bacteria towards the periphery and a favorable multiplication there. From the experiments on *E.oli* ^7^ and *Salmonella*^11^ different strategies for biasing the bacterial random walk, what we call *minmax* and *min* tumble rates were inferred. To make the discussion complete, we studied four different strategies listed in the methods section - the tumble rates towards and away from nutrients being affected or unaffected. In Figure 5B (*max* tumble rate) though there is a formation of ring but the nutrient uptake and cell division are lesser than that of the strategy corresponding to the *minmax* tumble rate (Figure 6). In Figures 5C (*min* tumble rate) and 5D(*fixed* tumble rate), however the bacteria can also drift away from the nutrient source with a higher chance thereby reducing the overall nutrient gain and population growth and not forming the rings. The quantitative details are shown in Figure 6, where the number of bacteria and nutrient absorbed are shown in Figure 6A and Figure 6B respectively. The number of bacteria increased in steps in all these calculations. In order to understand the origin of this step like increase, we perfomed a seperate simulation where the initial bacterial population did not just have a single size (*m_D_*) but rather a mixed age population with a distribution of bacterial sizes between just divided (*m_D_*) to ready to divide (2*m_D_*). This resulted in in a smooth increase in bacterial population for *minmax* tumble rate. The maximum rate of bacterial growth occurs only with the *minmax* chemotactic strategy. While a clear signature of chemotactic advantage is seen, in this simulation of lab experimental conditions it must be noted that the concentration as well as the amount of nutrient are very high compared to most realistic conditions.

**Figure 5:**
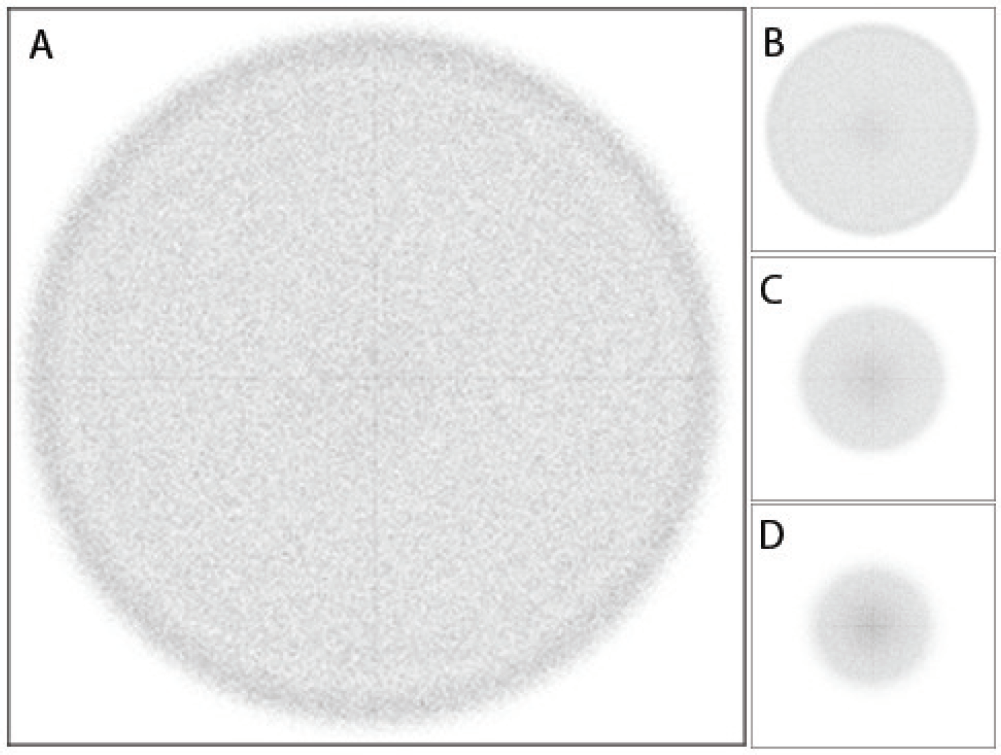
Simulation of an experiment where a bacterial colony is placed at the center of a petridish (40 mm side) with an initial uniform nutrient concentration of 100 *µ*M. The calculations were performed to capture 8 hours of laboratory conditions using four different tumble rate strategies A. *minmax* tumble rate, B. *max* tumble rate, C. *min* tumble rate, and D. *fixed* tumble rate. When walk is biased in both directions, the experimentally observed rings appear within the 8 hours of simulation. Images are given at 6 hours of experiment time

**Figure 6:**
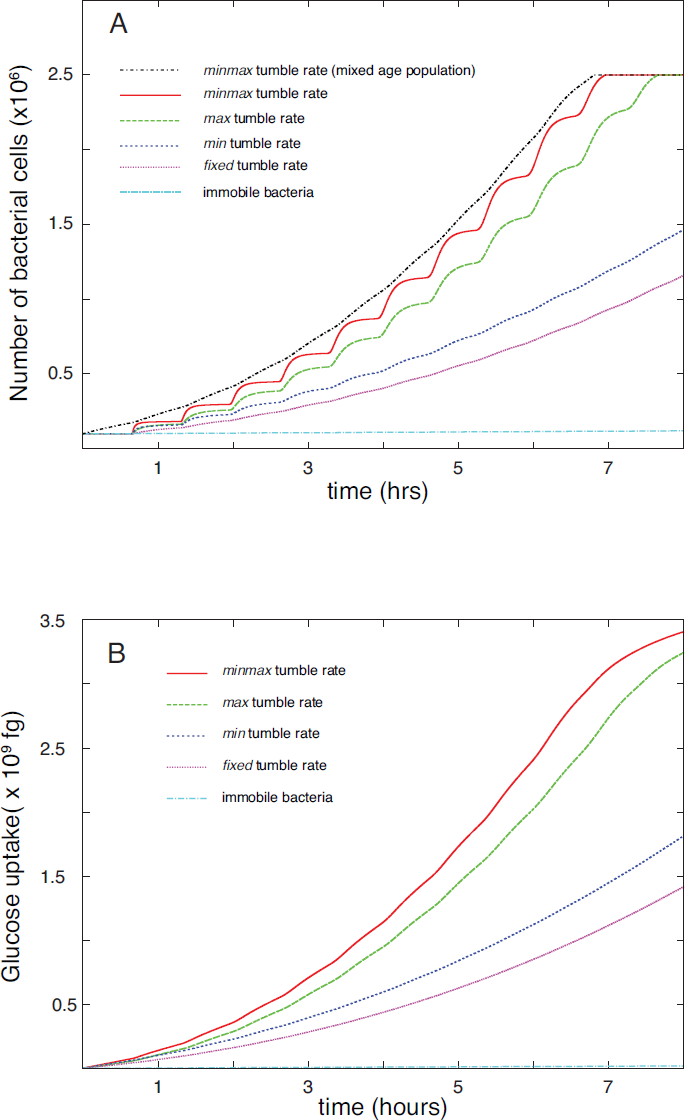
Quantitative results from the simulations replicating the petridish experiments: (A) cell division and (B) glucose uptake over 8 hours depending on the random walk strategy. The step-like behavior in most of the nutrient uptake curves is because all bacteria started from same age (size). *minmax* tumble rate strategy was repeated starting with the bacterial colony having a size distribution from just divided to ready to divide and it shows a smoothly increasing cell count.

#### Effect of absorption kinetics on bacterial growth rate

The absorption kinetics depends on the type of the nutrient and kinds of absorption pathways which are active, etc. We restrict our analysis to the absorption of glucose via Michaelis-Menten kinetics. In the literature all nutrient molecules colliding with bacteria are assumed to be perfectly absorbed. While this perfect absorption may be a good model under nutrient limited conditions but over predicts the cell division, absorption and when the nutrient is sufficiently available in nutrient rich experimental conditions. The exponential growth phase using the two types of kinetics in our model predicts the time for population doubling to be 90 min (Michaelis-Menten kinetics) and 6.5 min (complete absorption) respectively (Figure S16). The doubling times of 6.5 minute is too fast to be true, and perfect absorption models should be used with caution.

#### Effect of initial population on swarm ring formation

Swarm ring formation depends on the nutrient availability as well as its depletion at the center by the bacteria. The size of the initial population clearly has a significant role on when the rings form. We have simulated eight hours of laboratory experiment in a petridish with 100 *μ*M initial nutrient concentration with three different population sizes - 10^3^, 10^4^ and 10^5^. All the three initial population in this range showed ring formation for *minmax* tumble rate strategy (Figure S15), but population with smaller size showed smaller rings and formation of such rings took larger time. Specifically, with all the microscopic details accounted for by our model, the 10^5^ initial population showed a ring which extended upto the boundaries of the dish within a simulation corresponding to 8 hours of experiment. The appearance of the observed macroscopic pattern within experiment time gives us confidence that the microsimulation starting with all microscopic details can model the population level observations.

#### Growth advantage of chemotactic bacteria

In order to study the relative growth advantage of bacteria having chemotaxis mechanism, we simulate mixed populations containing a chemotactic (*minmax* tumble rate) and non-chemotactic (*fixed* tumble rate) in the same conditions as in swarm ring formation noted earlier. The total number of bacteria at start was kept at 10^5^, and the chemotactic fraction was varied as: 1%, 10%, 30% and 50%. Even when the initial fraction of chemotactic bacteria in the population was 1% within 8 hours of experiment time the chemotactic population surpasses its non-chemotactic counterpart. The X-Y projection of the position of chemotactic and non-chemotactic bacteria at different time point (Figure 7) reveals the preferential movement of chemotactic bacteria away from nutrient depleted regions leads to greater population growth compared to the non-chemotactic bacteria (Figure 8). The chemotactic population shows even faster growth rate compared to the non-chemotactic population when we start with 10%,30% and 50% chemotactic bacteria in the population (Figure S18) Comparing the nutrient gain between chemotactic and random walk mechanisms single *E. coli* and the colony under different nutrient conditions, we find that chemotaxi movement with its inherent noisy sensing limits helps in going from nutrient rich to nutrient richer conditions rather than from nutrient poor to nutrient rich conditions. The detailed limits of advantages of chemotaxi will be explored later.

**Figure 7:**
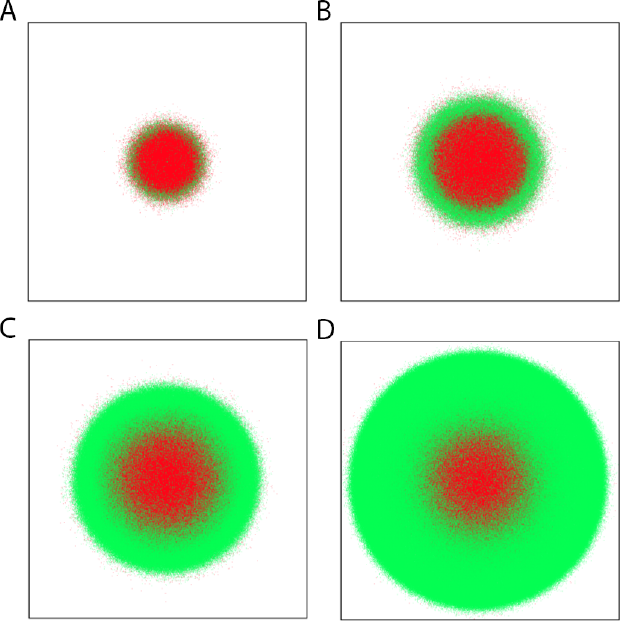
X-Y projection of the position of chemotactic (green) and non-chemotactic (red) bacterial populations after (A) 2 (B) 4 (C) 6 and (D) 8 hours from start. The initial fraction of chemotactic bacteria is 1% and it is placed a petridish of 40*mm* × 40 *mm*, represented as the square in each of the pictures

**Figure 8:**
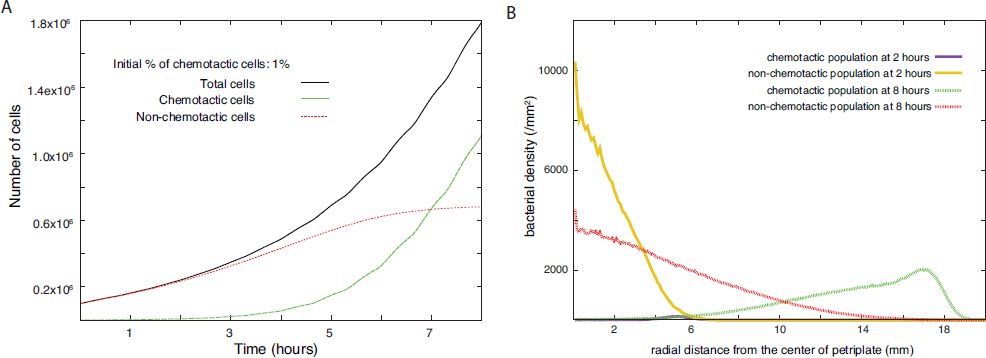
Quantitative description of the growth of a mixed population starting with an initial 1% chemotactic and 99% non-chemotactic bacteria: (A) Over an 8-hour period the chemotactic bacterial population multiplies over a hundred times faster and grows more than the non-chemotactic one (B) The radial distribution of the population 2 and 8 hours from the start shows the chemotactic and non-chemotactic bacteria preferring distributions around the periphery and the centre.

## Conclusion

A microsimulation method which can track every single bacterium in a large population has been developed. The simulation seamlessly integrates several interesting details at a single bacterium level to reach predictions for a population. Using the detailed model we are able to make comparisons of the nutrient gain by bacterium in different conditions. The model offers the flexibility of adding further assumptions at a single bacterium level and make comparisons with experimental studies.

## Supplementary information

### Diffusion equation

The nutrient molecules in the medium gets either di use or get depleted owing to uptake by the bacterial population following,

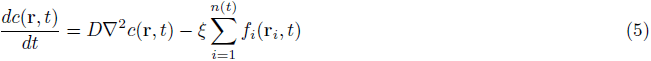

*c*(**r**; *t*) is the chemical concentration at position **r** at time t. For our simulation we need to discretize space and time to solve the diffusion equation using finite difference method. We discretize the space using dx=dy=dz=25*μ*m and time using 
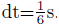
 We also use the diffusivity D=600 *μm^2^/s* similar to the diffusivity of glucose^49 50^ and 
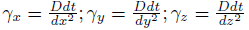
.
The indices for space, time discretization are given by 
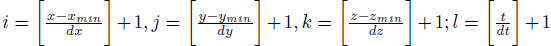
 in our simulations, where *min* corresponds to minimum values of an entity and [q] corresponds to the integer part of a real quantity q. Let *f(i, j, k, l)* be the amount of glucose consumed between time t and t + dt and inside grid (*i, j, k*), 
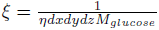
is the conversion factor from mass to concentration *η* = 602.3*μM^-1^μM^-3^* and *M_glucose_* is the molar mass of glucose. We solve the diffusion equation using the following finite difference method,

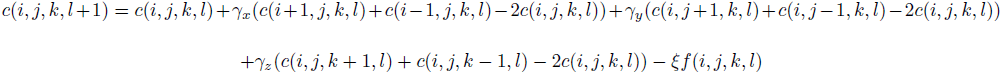

with zero-flux boundary condition as,

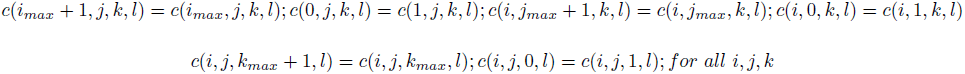

where, *c(i, j, k, l)* is the chemical concentration at time *t = l × dt* in the *(i, j, k)^th^* grid.

### Calculating maximum uptake rate for Michaelis-Menten kinetics

The excess uptake rate of glucose for 1g dry weight of *E. coli* per hour in presence of other sugars is ~1.7g^43^. Based on the above data we assume the maximum uptake rate of glucose in absence of any other sugar is ~2*g* per gram of dry weight of *E. coli* per hour. Further we know the average mass of an *E. coli* is 1 pg and almost 2/3 of it is water^44^ so, 1 g of dry weight of *E. coli* contains ~3 × 10^12^ cells. So a single *E. coli* can uptake with a maximum rate of 
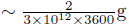
 glucose/second or ~0.19 fg of glucose per second.

### Experiment with a nutrient rich blob

We simulated a bacterial population using microsimulation approach starting with 10^5^ bacteria starting at a distance of about 8500*µm* from the glucose rich source of initial concentration 10,000 *µ*M and of radius 1250 *µm* centered at the origin. We quantified the cell division and nutrient uptake for *minmax, max, min*, fixed tumble rate and immobile bacterial population. The cell division(Figure S10) and nutrient uptake(Figure S11) is most advantageous for *minmax* tumble rate followed by *max, min*, and *fixed* tumble rate with immobile bacteria being the most disadvantageous. The simulation time corresponds to 8 hours of real time for each strategy where details of noisy nutrient sensing, memory dependent bias in run and tumble motion, glucose uptake using Michaelis-Menten kinetics and cell division have all been considered at a single bacterium level.

**Figure S9:**
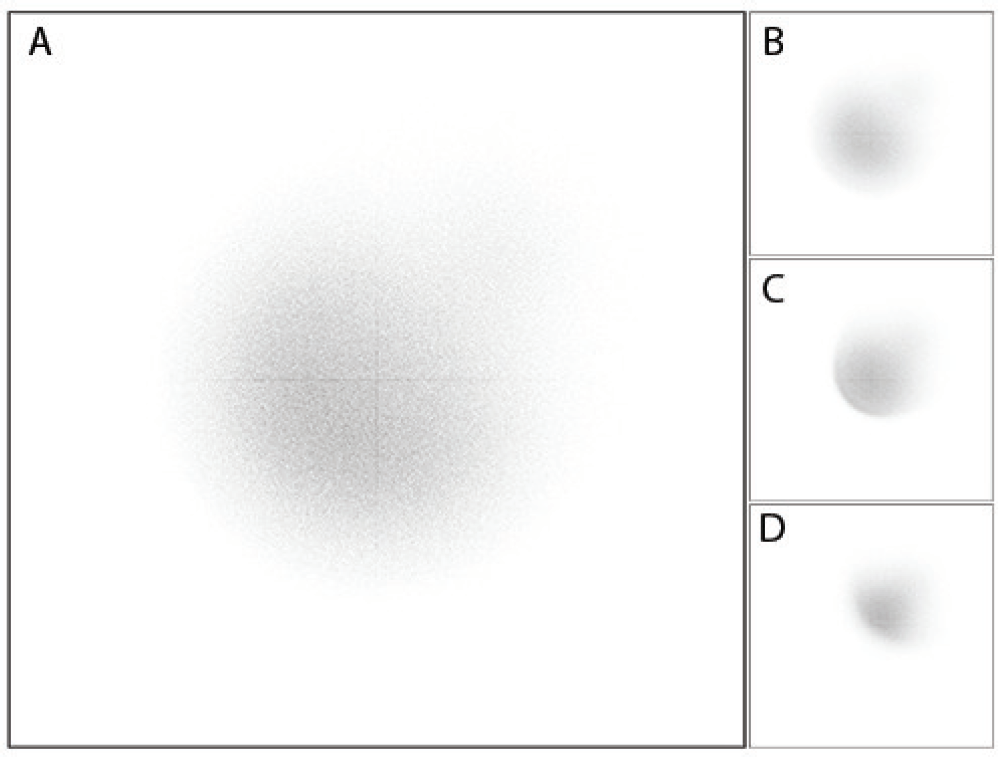
Population of bacteria after 6 hours starting at a distance of ~8500 *µm* with glucose rich source at the center. Each square has sides of 4cm. (A) *minmax* tumble rate (B)*max* tumble rate (C)*min* tumble rate (D) *fixed* tumble rate

**Figure S10:**
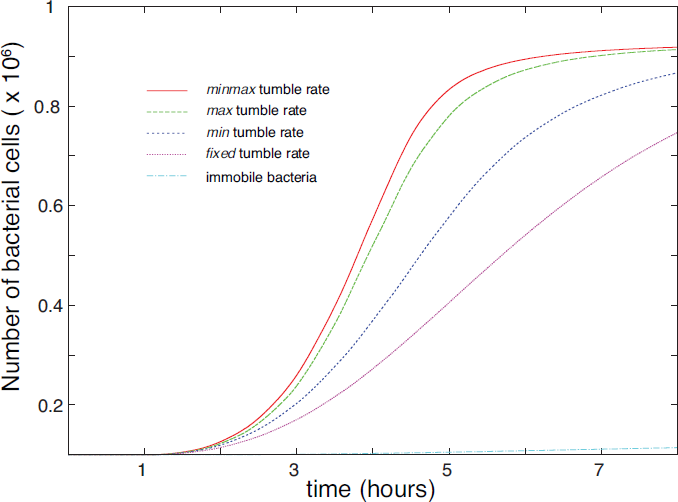
Cell division in 8 hours, starting with 10^5^ cells at (6000 *μm*, 6000 *μm*, 0) with the glucose rich source centered at the origin.

**Figure S11:**
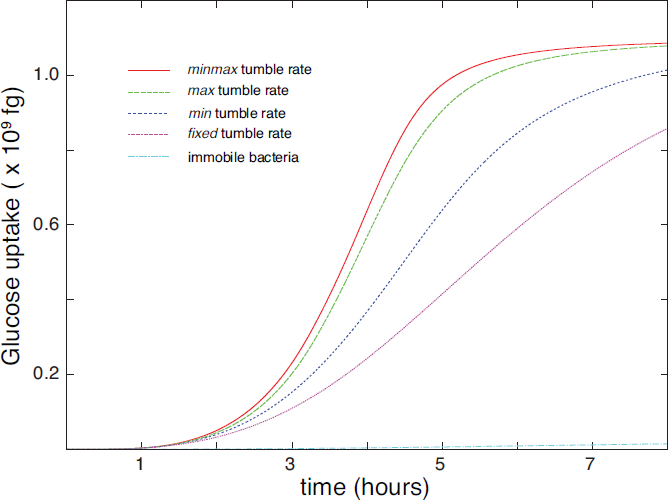
Glucose uptake(femto-gram) in 8 hours, starting with 10^5^ cells at (6000 *μm*, 6000 *μm*, 0) with the glucose rich source centered at the origin.

### Phytoplankton lysis event

We perform microsimulation on a population of bacteria starting with 10^5^ cells from a distance of 900*μm* from the glucose rich source of initial concentration of 1000*μM* having 100*μm* radius at the centre of a box of dimensions 4*mm* × 4 *mm* × 4*mm*. This mimics the natural event of phytoplankton burst in the ocean with a background glucose concentration of 40*μM*. Simulating for the various strategies reveals that under the given circumstances chemotaxis does not over any significant nutrient uptake advantage.

**Figure S12:**
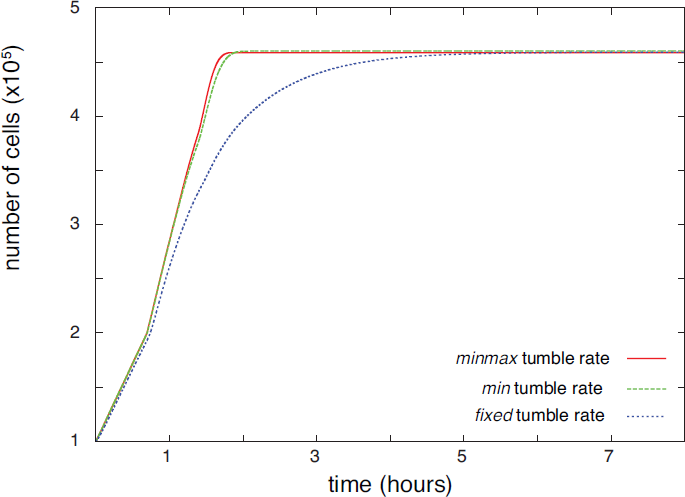
Cell division in 8 hours of phytoplankton lysis event starting with 10^5^ cells at (500*μm*, 500*μm*, 500*μm*). The step like characteristic is owing to starting with same size of each bacterium in the population.

**Figure S13:**
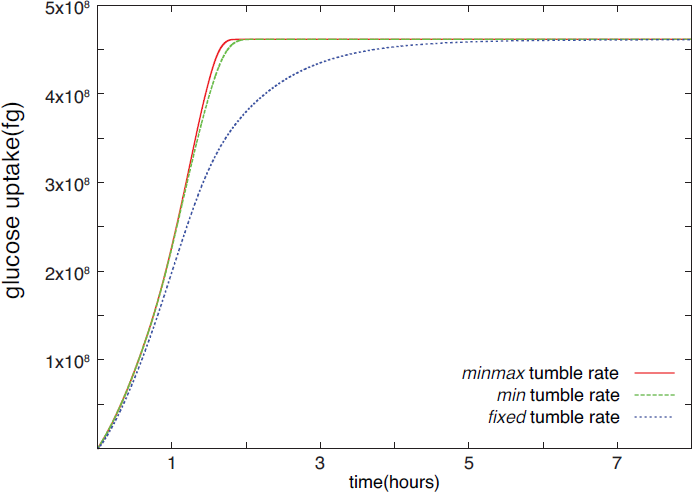
Glucose uptake(femto-grams) in 8 hours of phytoplankton lysis event starting with 105 cells at (500*μm*, 500*μm*, 500*μm*)

### Randomizing initial size

The bumpy nature of the cell division plot corresponding to the ring formation experiment is owing to the fact that we start with all bacteria having same size, hence they also performs cell division synchronously. This synchronous cell division vanishes(Figure 6) when we randomize the initial size of the population while forming the rings(Figure S14).

**Figure S14:**
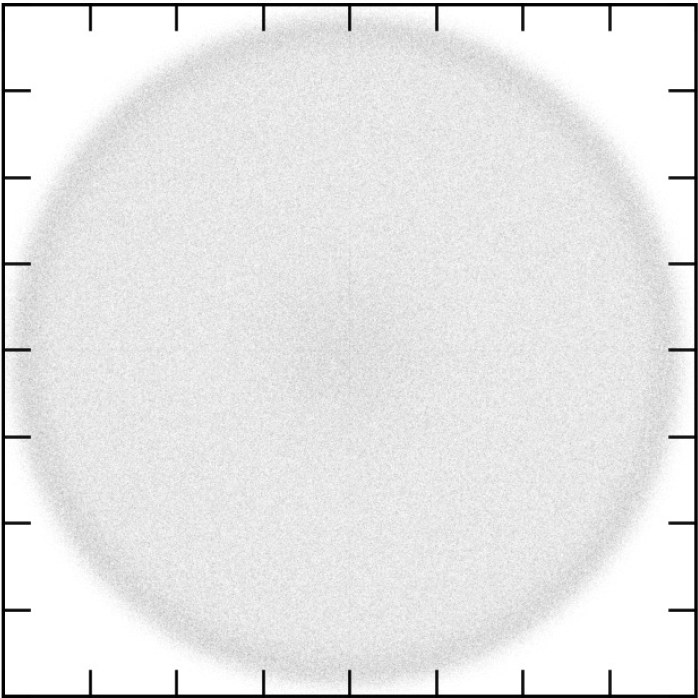
Randomizing initial size of each bacterium in the population did not affect ring formation (*minmax* tumble rate).

### Effect of Size: Ring formation after 6 hours

We perform the microsimulation for swarm ring formation with chemotactic bacteria (*minmax* tumble strategy) for 8 hours of laboratory experiment time with three starting population sizes (*n*_0_ = 10^3^, 10^4^, 10^5^). All population forms rings, but the sizes of the ring decreases with initial size of the population.

**Figure S15:**
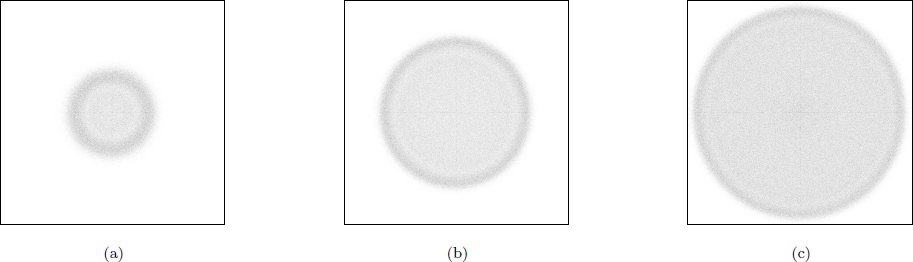
Swarm ring formation for initial colony sizes of (a) n_0_ = 10^3^, (b) n_0_ = 10^4^, (c) n_0_ = 10^5^ bacteria. The size of the square in all cases is 40 *mm* × 40 *mm*

### Perfect Absorption vs. Michaelis Menten Kinetics

We compare the glucose uptake and growth of *E. coli* population using perfect absorption and Michaelis-Menten kinetics. Perfect absorption corresponds to uptake of all glucose molecules that collides with the bacteria. Cell division takes place based on mass doubling (absorbing 900 fg of glucose). The calculated doubling times for uptake using Michaelis-Menten kinetics and perfect absorption was found to be 90 minutes and 6.5 minutes respectively. The latter doubling time is too fast for practical purposes.

**Figure S16:**
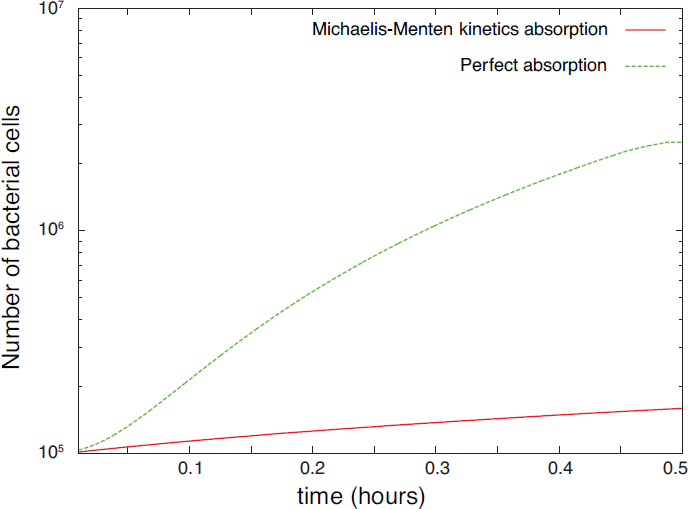
Comparison of cell division for different ways of glucose uptake for minmax tumble rate in swarm ring experiment.

**Figure S17:**
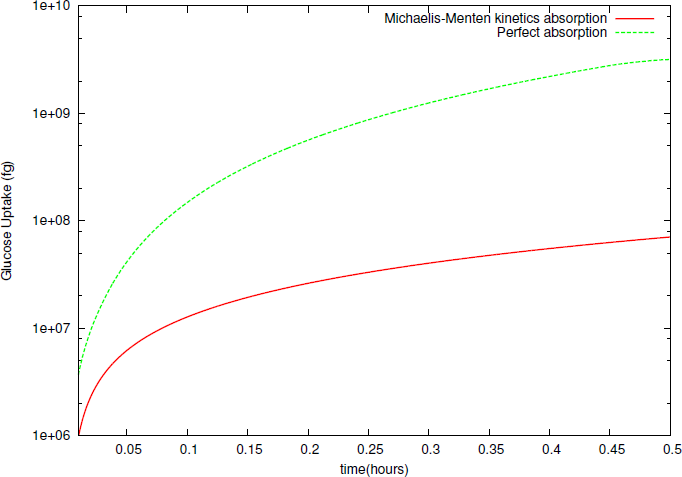
Comparison of glucose uptake using different ways for *minmax* tumble rate in swarm ring experiment.

### Growth advantage of chemotactic bacteria

**Figure S18:**
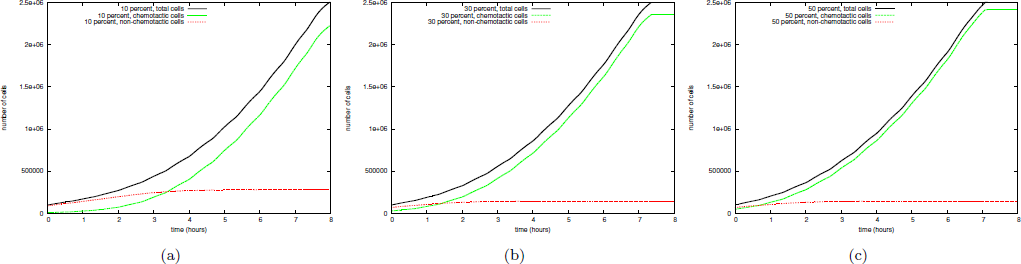
Growth advantage of chemotactic (green line) over non-chemotactic (red line) bacteria over 8 hours experiment time starting with (a) 10%, (b) 30%, (c) 50% chemotactic population. Black line represents the total population.

